# Microfluidic Live-Imaging technology to perform research activities in 3D models

**DOI:** 10.1101/2021.03.08.434339

**Authors:** Capuzzo Arnaud Martino, Daniele Vigo

## Abstract

One of the most surprising differences observed when comparing cell cultures in 2D and 3D is morphological dissimilarity and their evolution over time. Cells grown in a monolayer tend to flatten in the lower part of the plate adhering to and spreading in the horizontal plane without expanding in the vertical dimension. The result is that cells grown in 2D have a forced apex-basal polarity. 3D cultures support co-cultivation and crosstalking between multiple cell types, which regulate development and formation in the in vivo counterpart. 3D models culture, with or without a scaffold matrix, can exhibit more *in vivo-like* morphology and physiology. 3D cultures recapitulate relevant physiological cellular processes, transforming into unique platforms for drug screening. To support and guarantee the functional maintenance of a 3D structure, one must consider the structures and dynamics of regulatory networks, increasingly studied with live-imaging microscopy. However, commercially available technologies that can be used for current laboratory needs are limited, although there is a need to facilitate the acquisition of cellular kinetics with a high spatial and temporal resolution, to elevate visual performance and consequently that of experimentation. The CELLviewer is a newly conceived and developed multi-technology instrumentation, combining and synchronizing the work of different scientific disciplines. This work aims to test the system with two models: the first model is a single *Jurkat* cell while the second is an *MCF-7* spheroid. After having grown both models, the two models used are loaded into the microfluidic cartridge for each experiment and recorded in time-lapse for a total of 4 hours. After adaptive autofocus, when sliding inside the cartridge chamber, the samples used are tracked under the action of the optics and the 3D rotation was experimentally successfully obtained. A cell viability assessment was then used using the MitoGreen dye, a fluorescence marker selectively permeable to live cells. The ImageJ software was used to: calculate the model diameter, create fluorescence intensity graphs along a straight line passing through the cell, visualize the spatial fluorescence intensity distribution in 3D.

## INTRODUCTION

Cultivating cells and making cell models, in a biotechnological or biomedical laboratory, under controlled and favorable conditions, is essential for scientific research with aims in different fields of study: the use of stem cells in regenerative medicine, from the discovery of targets and cancer drugs, the production of therapeutic proteins and finally the modeling and characterization of diseases [1]. *In vitro* cell cultures can be isolated from normal or diseased tissues, can be grown as adherent monolayers or in suspension, and can be arranged to summarize some of their functions, in two or three dimensions [2], [3].

The two-dimensional (2D) approach involves cells that proliferate on flat substrates, resulting in a monolayer cell far from the type of cells that grow *in vivo.* They are devoid of cell-cell and cell-matrix interactions in the microenvironment[4].

The traditional 2D culture systems growth alone or in co-culture on plates, in which experiments supported by *in vitro* imaging are conducted for different functional, pharmacological, toxicological, and even clinical applications; they have long been widely used and already known for the nature of their cost and high repeatability. However, 2D culture systems cannot reach a stage of 3D organization equal to *in vivo*, due to the disadvantages associated with the lack of specific tissue architecture, mechanical-biochemical signals, cell-cells and Extracellular Matrix (ECM) [5]–[7].

One issue with conventional 2D cell culture systems is the inadequate quality and quantity of ECM, which is fundamental to the support of the structure by facilitating communication between the different cell populations embedded in the matrix by imparting mechanical properties to the tissues[8]. Cells in 2D culture are not surrounded by ECM and therefore are different from the structure of an in vivo cell system, as they cannot: migrate, polarize, differentiate in response to[9]–[12]. Despite their proven value in biomedical research, 2D models cannot support differentiated and cell-specific functions in tissues or accurately predict in vivo tissue functions and drug and biological modulator activities [13]–[16]. These limitations have led to a growing interest in the development of more complex models, such as those that incorporate multiple cell types or involve cell modeling, and in three-dimensional (3D) models, which better represent the spatial and chemical complexity of living tissues[17]–[19]. This lack has led to the development of three-dimensional (3D) cell culture models to improve *in vitro* research systems, more accurately recapitulating the *in vivo* state in which: cell morphology, interactions and specific architecture most represent that of native tissues[20].

Three-dimensional (3D) cell culture better replicates the physiological interactions between cells and their environment than traditional two-dimensional (2D) cell culture cells. Particular types of cell lines have the ability to self-aggregate into 3D aggregates called cell spheroids when grown in suspension [21].

The terms spheroid and organoid are commonly used when addressing the topic of 3D cell culture, but this terminology has been used interchangeably. There are several distinct differences between them for equal merit of definition, implementation and utility as an advanced model of scientific research [20], [22], [23].

### 3D Models

#### Definitions and Current Terminology

Spheroids and organoids both refer to 3D culture models that have specific organization and architecture, not achievable through 2D monolayer cell cultures. There is currently no universally well-defined nomenclature for these models and in the literature.

The term spheroid, first coined in the 1970s, was noted that some Chinese hamster lung cells when dissociated were capable of forming almost perfectly spherical suspended cell aggregates grown in rotating flasks[20]. Since that time, spheroids have been generated from many types of primary cells and cell lines in a multitude of methods such as hanging drop, low cell attachment plates, micropatterned surfaces or rotating bioreactors[3], [20], [24]–[26].

The term spheroid is currently used in research where cancer cells constitute the invaluable multicellular model of tumor spheroid (MCTS) for studying solid tumor biology[20].

The spheroid is used to describe tumor models in which the 3D structure has been formed from standard immortalized cell lines traditionally grown in 2D. Spheroid cultures can also be generated from primary tumor cells [27].

The MCTS spheroid is characterized by having a subdivision into regions. Starting from the most external there is the proliferation of cells, an internal zone in which quiescent cells reside and finally the central zone composed of a necrotic core[20], [24]. This arrangement mimics the cellular heterogeneity observed in solid tumors[28].

The cell arrangement within the MCTS emulates cell morphology, proliferation, oxygenation, nutrient uptake, waste excretion, and drug uptake *in vivo*[20], [29].

The embryoid bodies are 3D stem aggregates capable of giving rise to cells that form the 3 germ layers and in turn differentiate into aggregates[20]. When incorporated into Matrigel without the addition of specific growth factors, the aggregates form buds and further develop into discrete regions[30].

Embryoid bodies, mammospheres, hepatospheres and neurospheres are the most widely used spheroid models as a MCTS from cancer cells. They are used as models of avascular cancer to obtain information on cancer invasion and therapeutic cancer screening of metastases [20], [30]. The subdivision of the population that makes up the spheroid heterogeneously, between those in proliferation and not, exposes the spheroid model to an oxygen gradient that emulates the physiochemical gradients found in solid tumors[28]. Some advantages of spheroidal 3D models are that they are relatively easy and quick to model, versatile depending on the study under consideration, and cheap enough to tackle high-throughput screening[31].

The organoid was previously defined as an aggregation of cells that contained differentiated cells with some tissue-like structures[32]. The description currently associated with the term organoid is that of a 3D structure grown from organ stem cells or progenitor cells composed of organ-specific cell types that self-organize through cell selection and a space-limited lineage that mimics at least one organ function[20], [33], [34].

Organoid describes 3D biological structures grown *in vitro* that resemble their *in vivo* counterpart in architecture and function[23]. Characteristics of organoids include self-organization, multicellularity, and functionality[20]. They then recap the microanatomy of the organs with differentiated cell types specific to each organ[35].

Organoids can be mainly classified into two macro categories: those deriving from adult tissues and those deriving from pluripotent stem cells[3], [20], [24], [36], [37]. Organoids contain multiple cell types organized into structures that resemble the organ of interest and exhibit some of the functions of the study target organ[20].In general, a higher order of self-assembly is found in organoids than in spheroidal cultures, which are more simplistic, with little or no tissue structure characteristics[23]. The formation of organoids depends more on the presence of biological or synthetic matrices[29]. The interaction between the cell and the ECM is essential for survival, proliferation, differentiation and migration[25]. The ECM also provides a physical structure on which cells can move and grow in 3D[38]. Spheroid cultures are less dependent on matrices for their formation, capable of forming in both scaffold-free and scaffold-based conditions[20].

#### Advances in cell cultures

The development of 3D culture is closely related to understanding the cellular microenvironment and the importance of the ECM as a regulator of morphogenesis and tissue function[39]. The ECM is made up of insoluble collagen proteins and fibers. Collagen provides rigid structures and physical support, while ECM proteins, including fibronectin, laminin and integrins, are involved in signaling processes [24]. The basement membranes, or lamina propria, provide epithelial and endothelial cells with structural and organizational stability, act as a barrier, transduce signals and other functions[40].

Matrigel, isolated from chondrosarcoma in 1977, the first basement membrane extract, has been repeatedly used for 3D cell cultures and for suspension methods to form spheroids [41].

Following Matrigel and the characterization of ECM components, 3D culture approaches incorporate the use of matrices, animal-derived hydrogels or synthetic hydrogel networks with ECM components. Providing physical support and biochemical signals that recapitulate cell-matrix interactions. Thus incorporating biological aspects of morphogenesis, homeostasis and recapitulating spatial signals that appear to be absent in 2D cultures [29], [41].

An advantage of cell growth in 3D cultures is that they are reflected more accurately *in vitro* than cells growing in monolayer cultures. This is because cells do not normally grow or interact in isolation; rather, they form interactions with other cells and the microenvironment that surrounds them [42]. By affecting gene expression, the distribution of proteins such as receptors, transcription factors or cell cycle regulators [43].

Cell-cell interactions in 3D cultures involve cells of the same type, as can occur in spheroids grown from a tumor cell line. Communications can involve multiple cell types when co-cultured with other cells [20].

Cultures of tumor spheroids exhibit tumor-like cytoarchitecture and contain metabolic gradients of gas and nutrients. As the distance from the periphery increases, the availability of oxygen and glucose decreases, reflecting the state of hypoxia and nutrient deprivation, characteristic of neoplasms, which influence the reactivity and efficacy of the drug. Spheroid patterns also recapitulate lactate accumulation in tumors, which modulates tumor immunogenicity, reducing immune receptor expression and inhibiting effector cell action. These models also summarize the proliferative gradients of tumors: an outer layer of proliferating cells surrounds a layer of non-dividing but viable cells, defined as the drug-resistant dormant niche. The increase in cell death by apoptosis and necrosis leads to the accumulation of a necrotic core, which accumulates in the center of the spheroid [25], [33], [44].

The biological features of 3D tumor models, such as modulation of immune receptor expression and accumulation of lactic acid, mimic the *in vitro* processes of tumor evasion from the immune system. They can therefore be used to evaluate immunotherapeutic approaches [29].

3D cultures recapitulate relevant physiological cellular processes, transforming into unique platforms for drug screening. 3D culture models replicate many factors affecting anticancer drug activity *in vivo*, proliferation gradients, cell-cell interactions and ECM-cell signaling, non-uniform exposure of cancer cells to oxygen and nutrients, microenvironment [45], [46].

Current strategies incorporate recent advances in stem cell biology, following ESC from mouse blastocysts in 1981 and humans in 1989, the introduction of somatic cell pluripotency factors generating iPSCs in 2006 [47].

The multitude of advances in stem cell biology have enabled the formation of 3D organoid cultures [48].

Organoid culture technology has made huge strides in the last decade leading to the development of models, which mimic different organs including: intestine, stomach, pancreas, colon, liver and many more [20], [36], [49]–[52].

Organoids grown in 3D represent *in vitro* organ development, structural properties and organ functionality *in vivo*. They are used to model the development and physiology of organs in pathological states, and the fate of an organoid can be modulated through genetic editing, as occurs in genetic disorders and infectious diseases[20].

Organoids also have multiple applications in drug discovery and toxicity assays[53]. Disease modeling in organoids may involve pathogen infections or the targeted introduction of pathogenic mutations including inherited mutations or cancer development and progression[54].

Although the use of 3D cultures recapitulate unique aspects of cell function, current 3D cultures represent reductionist models of the *in vivo* counterpart. Unfortunately, 3D models lack several aspects of organ biology and pathological processes such as the presence of surrounding tissue types, innervation, vascularity, the presence of immune cells and tumor stroma[20], [55], [56].

#### Co-culture system

Co-culture systems have long been used to study the interactions between cell population and are fundamental to cell–cell interaction studies. Such systems have become of particular interest to biologists for the study and design of complex multicellular 3D systems [57], [58]. Co-culture allows a variety of cell types to be cultured together to examine the effect of one culture system on another, useful when examining the effect of one type of tissue, region, or how a particular molecule secreted leads to changes in development or physiology[59], [57], [60], [61].

Co-culture is a cultivation set-up commonly adopted in 3D spheroid and organoid cultures, where two or more different cell populations are grown with some degree of contact with each other and the use of such a setup includes: study natural interactions between populations, improve crop success for certain populations or establish interactions between populations[62], [63].

2D co-culture is traditionally used to investigate the interactions between cancer cells and stromal cells in tumor tissues, but often fails to reproduce *in vivo* cell responses due to the highly artificial culture environment. Currently, 3D cell culture systems are being widely adopted in the fields of biological research, drug discovery and tissue regeneration[63], [64].

Compared to 2D monolayer cultures, cells in a 3D models culture, with or without a scaffold matrix, can exhibit more *in vivo-like* morphology and physiology such as proliferation rate and gene expression pattern[65]. The main advantages of 3D systems over 2D monolayers for co-culture studies include: 3D cultures enable the formation of spheroids or spatial cellular organization; Across the ECM and 3D cellular structure, gradients of nutrient, oxygen and growth factor can be created and play important roles in the tumor-stroma crosstalk; In 2D co-culture, different cell types are usually mixed and grow on the same monolayer. In contrast, 3D systems allow the co-cultured cells to be situated in different compartments or matrix layers [63].

3D cultures support co-cultivation and crosstalk of multiple cell types, which regulate development[66].

#### 3D models become fundamental for analyzing cellular relationships

Some 3D models provide great results in representing tissue structures in the physiological field compared to two-dimensional 2D cell culture[10], [19], [67]–[69]. The fabrics have a hierarchical structure that contains micro-architecture features that can be studied on many length scales. These include the subcellular/cellular scale (1–10 μm), which affects cellular function; the multicellular scale (10–100 μm), which determines the type and degree of intercellular interactions; and the tissue scale (100–1000 μm), which correspond[7], [70]. Deciphering population heterogeneities is a long-standing goal in cellular biology. At the level of single cells, such heterogeneities are usually observed at the genomic, transcriptomic or phenotypic levels[71]. In general, spheroids, self-organizing and heterogeneous cell aggregates up to 400–500 μm in size, are used for research, resulting from the suspension or adhesion on the single-cell jamb or co-culture of more than[72]. Spheroidal models have advantages derived from their geometry and the possibility of developing effects in co-culture and sustainability generally long-term, as they mimic optimal cell-cell and cell-ECM physiological interactions, reproducibility and the similarity in protein-gene expression profiles. The use of these models is not transferable to cell types, as 3D spheroids of these cells tend to disintegrate or take unpredictable forms[21]. To avoid unpredictable and not currently useful developments, various types of scaffolds are manufactured and applied tools that control the development and structuring of spheroids’ uniform dimensions [22].

The realization of an organoid, pseudo-organ, or neo-organ has in common some processes present in the various stages of development and formation of a living organism. This includes differentiation, proliferation, polarization, adhesion and precisely controlled apoptosis that combined with self-organization and multi-cellular pattern leads to the development of the various districts[73]. The Organoids or Tissues Organs are an *in vitro* 3D cell cluster derived from stem cells or progenitors and/or donors that spatially organize themselves in a similar way to their counterpart *in vivo*[74]. In the structuring and organizations of culture systems and especially of co-culture systems cells must maintain an adequate phenotype compatible with the external cellular environment and the duration of this phenomenon must be particularly protracted over time. For adhesion-dependent cells, interactions with the surrounding ECM and neighboring cells define the shape and organization of cells. One of the most surprising differences observed when comparing cell cultures in 2D and 3D is the morphological dissimilarity and their evolution over time. Cells grown in a monolayer tend to flatten on the bottom of the plate dish by adhering and spreading on the horizontal plane without expanding into the vertical dimension. The consequence is that cells grown in 2D have a forced apex-basal polarity. This polarity is probably relevant for certain cell types such as epithelial cells, but it is unnatural for most cells especially those of cubic or multifaceted type. The mesenchyme, if incorporated into a 3D ECM, take on a starry morphology and polarize only by bottom-up during migration[16], [75].

To support and guarantee the functional maintenance of a 3D structure, one must consider the structures and dynamics of regulatory networks, increasingly studied with live-imaging microscopy[76].

However, commercially available technologies that can be used for current laboratory needs are limited, although there is a need to facilitate the acquisition of cellular kinetics with a high spatial and temporal resolution, to elevate visual performance and consequently that of experimentation[10], [77]–[79].

#### Microfluidic live imaging

2D models in Petri dishes allow for collective cell simulation and behaviors related to disease modeling and understanding but the advent of laboratory and organ devices on a chip shows that information obtained from 2D cell cultures on plates differs significantly from results obtained in microfluidic environments as they reflect more biomimetic aspects [30].

2D culture imaging does not allow to fully appreciate the morphology of the cell population and the three-dimensionality of the sample, one of the reasons could be the unappreciated evolutionary changes. The use of imaging is an essential requirement for the study of the structural and functional morphology of the neo-organ, of its positioning / polarization and of cell differentiation, allowing the *in vitro* modeling of even the most complex organs [31]. Combining live imaging with the ability to retrieve individual cells of interest remains a technical challenge[80]–[82]. Combining imaging with precise cell retrieval is of particular interest when studying highly dynamic or transient, asynchronous, or heterogeneous cell biological and developmental processes[80]. Technological advances have enabled a broad array of live-cell imaging approaches to investigate cellular processes, many of which require optically demanding imaging modalities to achieve the necessary resolution and sensitivity [83]. A new technology based on newly developed microfluidics and imaging techniques can enable the management and identification of the phenotype, the biological activities of the present populations of the present populations without destroying the 3D of an organoid or derived in culture or co-culture of progenitor organs and/or donors who self-organize in space/time like the *in vivo*[74], [84], [85]. However complete lab-on-a-chip devices that can work with the automated procedure and allows to see the behavioral cities of cells or their alterations to support microfluidic system, have not been prevented in the literature usage[67].

Currently from what can be found in the bibliography, we have found that commercially the only equipment available to perform some specific protocols is CELLviewer[86], [87].

The CELLviewer is a newly conceived and developed multi-technology instrumentation, combining and synchronizing the work of different scientific disciplines in the field of management of both simple and complex 3D culture systems, allows to maintain in the most natural conditions possible the three-dimensional structure, following it over time through high-definition time-lapse microscopy.

## CELLviewer

### What is CELLviewer

The CELLviewer, Fig. 1., is a lab-on-a-chip for cells or neo-organs to be managed in the absence of adhesion, designed by CellDynamics, Bologna Italy. This multi-technological system is composed of a hardware tool and disposable parts: microfluidic chips used to insert the sample into them. The device’s specific capabilities include environmental control, automatic change of cultural media, the ability to insert individual cells or neo-organs, and perform an optical analysis in the light field, darkfield, and fluorescence microscopy. Besides, CELLviewer technology allows through programmable software and automated execution of custom protocols remotely. The CELLviewer is an innovative approach that allows time-lapse monitoring and imaging of suspended 3D models in a dynamic medium, as well as keeping biological units alive and mimicking the environment more naturally than other systems. The CELLviewer system keeps 3D models in suspension, counteracting the gravitational fall of cells, by generating extremely finely controlled micro-currents. In addition to being used to raise the falling sample for gravity, these micro-currents perform two other functions: they allow the biological sample to be moved into the three dimensions in a controlled and custom manner and allow the change of the administration of external stimuli such as drugs, fluorescent probes, growth factors or more. With high-resolution time-lapse fluorescence imaging, either of individual live cells or pseudo-organs has grown inside a suspension, observed, and subsequently placed in a disposable cartridge. The basic system is surrounded by a series of measures that allow you to maintain the environment for cellular life: a disposable cartridge avoids contamination between one analysis and the next; the channels in which we generate currents have dimensions compatible with both spheroids and single cells; The system of changing the culture soil and administering drugs allows an airtight, sterilized connection.

**Fig. 1.**
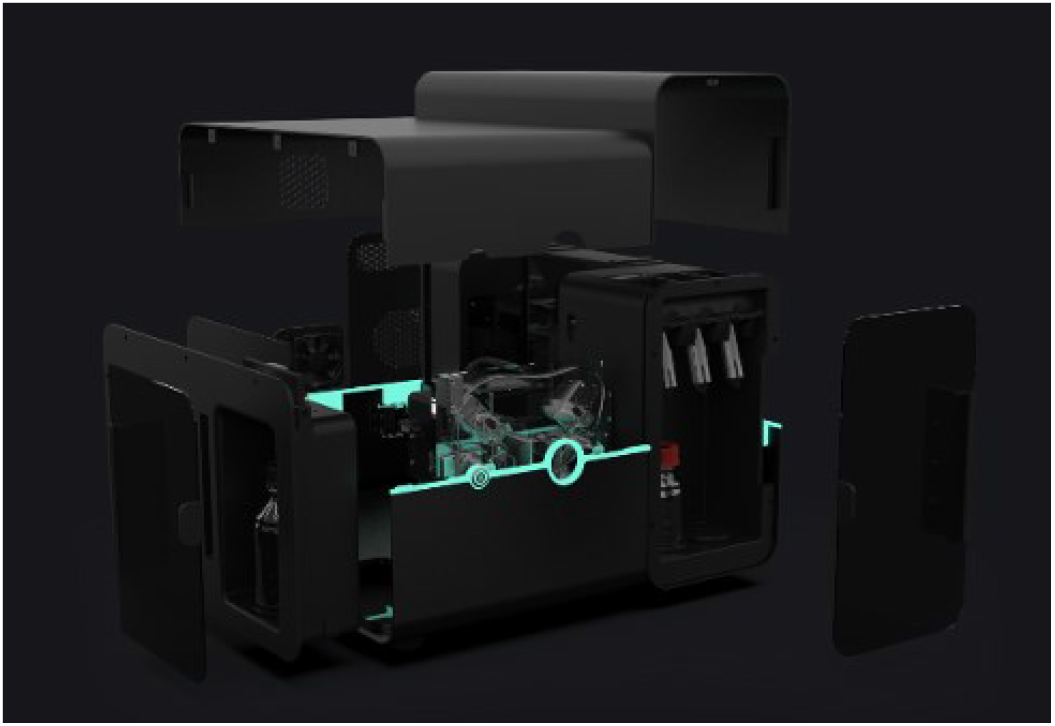
CELLviewer. The data obtained from CELLviewer allow you to observe and understand spontaneous or induced biological phenomena. To do this, to allow individual cells or various cell populations to be cultured and observed, the tool includes several optical and electronic components.

### CELLviewer components: Cartridge and Platform

The system mainly consists of a microfluidic integrated platform that combines different technologies, resulting in a hybrid system between an incubator for 3D cell culture and a wide-field microscope for time-lapse, live imaging. The platform is comprised of the hardware and the PC-embedded physical device, working on a single-use patented cartridge Fig. 2.

**Fig. 2.**
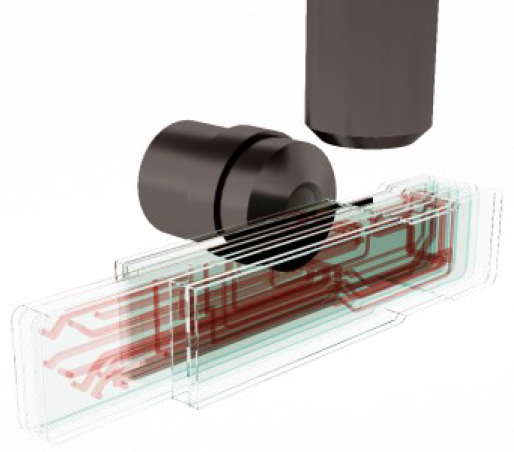
Cartridge. The central core of the microfluidic circuit includes three parallel analysis chambers with a rectangular shape and a volume of 3 μL each. Each analysis chamber rests on the lower side of the microfluidic circuit for sample load and medium culture replacement with drug solution or fresh swab. Four micrometric channels reach the upper side of every chamber to deliver buffer microflows that counteract gravity force, preserving the sample in suspension, while keeping it in focus position for time-lapse, live imaging.

The cartridge consists of multilayer plasma-bonded PDMS sheets and it is crossed by a complex network of microfluidic channels of different diameters (starting from 0.100 mm diameter to 1 mm diameter), covering a total volume of about 2 mL. It is hosted inside a warming cover in aluminum. Buffer microflows for sample management are generated employing four hydrothermal pumps.

The platform of CELLviewer is a hybrid technology that merges a fluidic module, an optical system, electronics, and software for the system management. The fluidic module is made by motorized rotative valves, selection valves, and pinch valves, and it is intended for the cartridge liquid management. The connection between the cartridge and the hardware system is realized via a motorized clamp connection.

The imaging setup has two cooperating light paths to acquire x-y-z coordinates of a single floating sample in the analysis chamber. Vertical optics has 2.5 X magnification (N.A. = 0.08) objective (Olympus Life Science) and a LD (low definition) camera, while time-lapse imaging is performed on horizontal path, equipped with 20 X (N.A = 0.45) and 40 X (N.A. = 0.60) objectives (Olympus Life Science), mounted on a motorized revolver for agile switching. HD (high definition) camera (Hamamatsu Photonics) acquires up to 30 frames/s in full resolution of 4.0 megapixels with a cooling element built-in, enabling long-term acquisitions. Illumination is based on LED stroboscopic light source that consistently reduces sample photodamage, with a white LED for sample positioning and bright-field imaging and 5 LED for multicolor Epi-fluorescence microscopy. The system is equipped with a 7-filter Pinkel penta-band set (Semrock Inc.), that is designed for imaging of a sample simultaneously labeled with DAPI, FITC, TRITC, Cy5, and Cy7.

### What we do with CELLviewer

To validate the ability of CELLviewer to grow biological samples, two experiments were carried out with different samples but in both the viability was assessed using MitoGreen-dye, a fluorescence marker selectively permeable to live cells. Cells can be stained with mitochondrial targeted dyes to understand cell status such as loss of apoptotic, high-energy or mitochondrial membrane potential (MMP). MitoGreen is a fluorescent mitochondrial dye that has shown green staining around to the nuclear region within the edges of the plasma membrane, which conceptually overlaps the mitochondrial region, contributing to the study of normal cell function in physiological states. MitoGreen-dye is a cell-permeating green fluorescent lipophilic dye that is selective for live cell mitochondria when used at low concentrations. Mitochondria are deeply involved in both cellular life and death mechanisms. In addition to serving as the main source of ATP production, mitochondria also function as a major buffer for calcium, which regulates the activities of enzymes. Furthermore, ROS generated by the electron transport chains of mitochondria can cause oxidative damage to cells. For these reasons, staining of mitochondria with fluorescent dyes, antibodies or naturally fluorescent molecules contributes to the study of their structure and function in normal physiological and pathophysiological states. The CELLviewer system enables high-content time-lapse fluorescence imaging of live cells grown in suspension within a disposable cartridge while dispensing drug solutions to the sample chamber to visualize their biological effects at the single cell level. Following this application notes [88], [89], single *Jurkat* cells and *MCF-7* spheroids are stained with MitoGreen and isolated within the disposable CELLviewer cartridge and finally time-lapsed in both Bright-field and GFP channels. MitoGreen is spectrally like FITC, making it excitable at 488 nm. MitoTracker dyes can be applied to measure the total mass of the mitochondria or to study the changes in the mass of the mitochondria following the desired treatments. Cell Signaling’s MitoTracker Green FM and ThermoFisher Scientific are examples of commercially available MitoTracker tests. MitoTracker Green dye stains mitochondria of living cells but does not depend on MMP MitoTracker Green is not compatible with fixation and the signal can be obtained at excitation and emission wavelengths of 490 and 516 nm respectively [79].

## EXPERIMENT AND RESULTS

### Single cell viability assessment

#### Materials

- *Jurkat* cells (ATCC)
- RPMI culture medium (Gibco, Life Technologies, Thermo Fisher Scientific)
- MitoGreen (PromoKine, PromoCell)
- CELLviewer imaging system
- CELLviewer 50 ml DOCK

#### Methods

The *Jurkat* cells were grown at 37° C and 5% CO2 in RPMI 1640 soil, supplemented with 2 mM of L-glutamine, 10% FBS, 100 units/mL of penicillin, and 100 mg/mL of streptomycin. Before the experiments, *Jurkat* were washed and suspended at final concentration of 5 × 10^5^ cells/ml in FBS culture soil at 5%.

Then, the sample was incubated for 20 minutes in the dark at 37 ° C with MitoGreen 200 mM (PromoKine, PromoCell). After incubation, the cells were centrifuged at 2000 rpm for 5 minutes to remove excess MitoGreen and resuspended in culture soil at 5% to the CELLviewer work concentration of 5 × 10^3^ cells/ml. The sample is then piped inside a 50ml Falcon tube closed with a 50ml CELLviewer DOCK. After isolation and fluid adaptive autofocus, CELLviewer automatically captures sample images in the Brightfield channel and GFP channel at 0.5 fps with 20X magnification.

After single *Jurkat* cell isolation in the microfluidic cartridge and fluid adaptive autofocus, CELLviewer automatically captures time-lapse imaged sample for 4 hours in the GFP channel, in Fig. 3., and Brightfield channel Fig. 5.

**Fig. 3.**
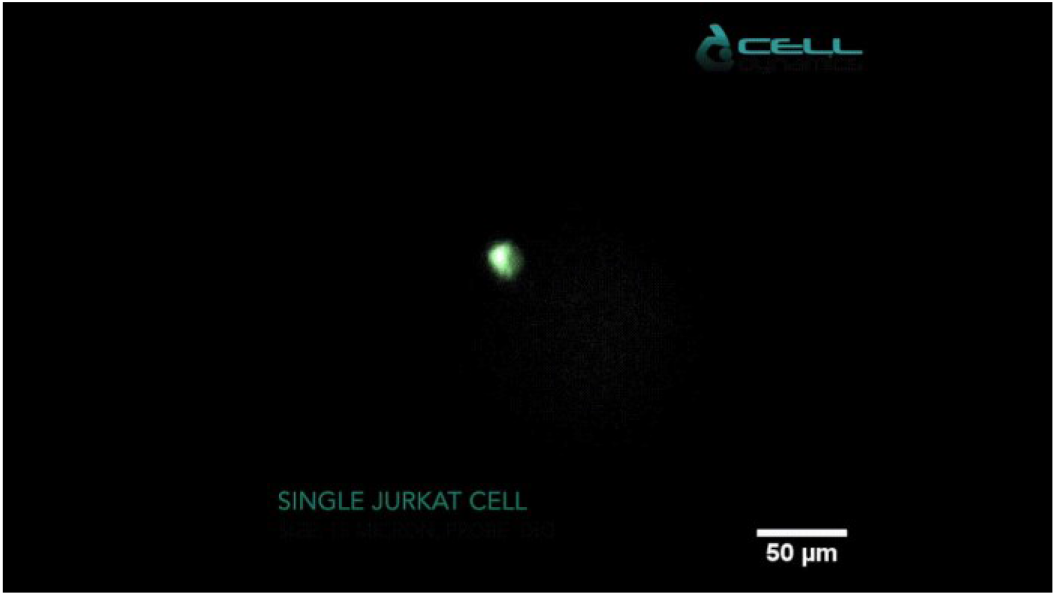
MitoGreen in live 3D single 3D cell. MitoGreen and resuspended in culture soil at 5% to the CELLviewer work concentration of 5 x 103 cells/ml. The sample is then piped inside a 50ml Falcon tube closed with a 50ml CELLviewer DOCK. It’s showed green staining around the nuclear region and within the edges of the plasma membrane. Scale bar: 50 μm.

ImageJ software was used for image analysis using the Measure function to: calculate the diameter of a single cell; Plot profile plugin to create fluorescence intensity graphics along a straight line that passes through the cell (Fig. 4); 3D surface plot plug-in to display in 3D the distribution of the intensity of spatial fluorescence. As we can see in Fig. 4. with the Plot profile and 3D surface plot, MitoGreen is a fluorescent mitochondrial dye that showed green staining around the nuclear region and within the edges of the plasma membrane, which conceptually overlaps the mitochondrial region, contributing to the study of normal cell function in physiological states.

**Fig. 4.**
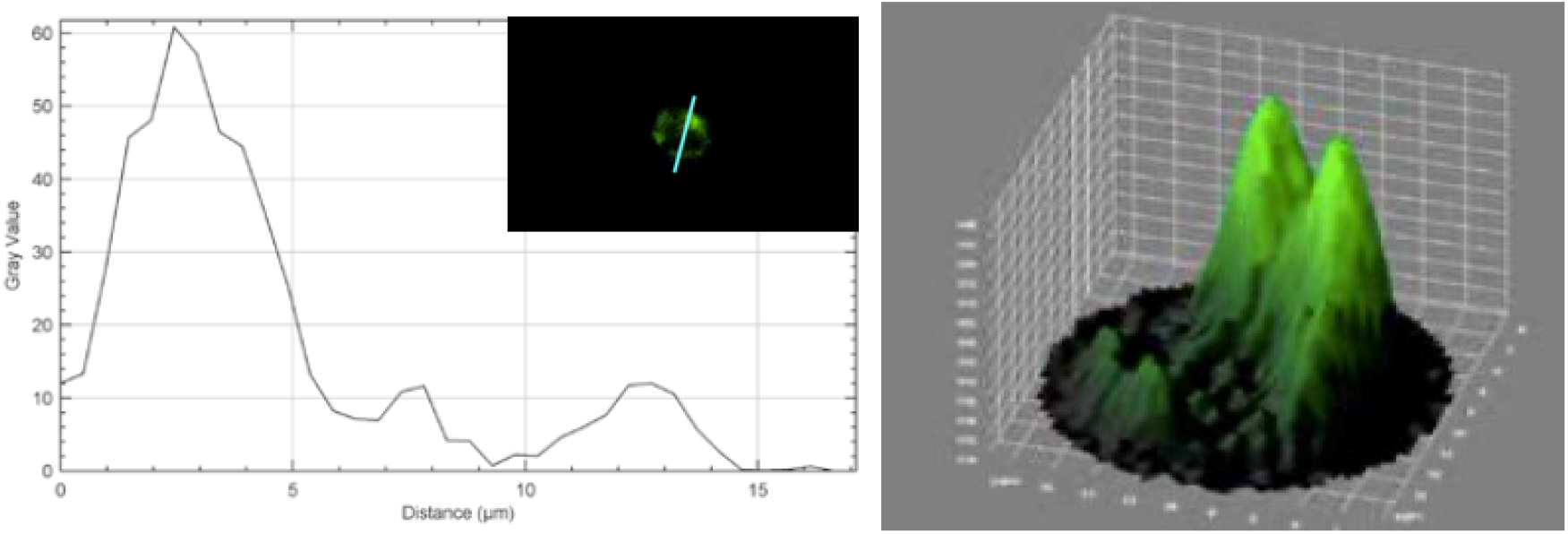
ImageJ software: Plot profile and 3D surface. Left: Plot profile plugin is used to create fluorescence intensity graphics along a straight line that passes through the cell. Right: 3D surface plot plug-in is used to display in 3D the distribution of the intensity of spatial fluorescence.

**Fig. 5.**
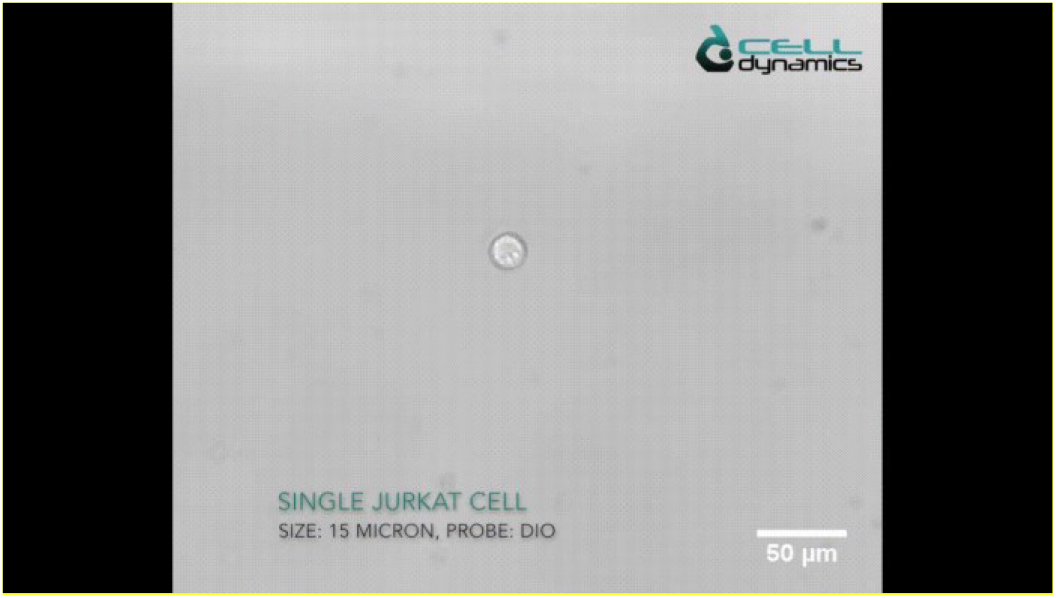
Single 3D live-cell. As is possible to see, when flowed inside the cartridge chamber, the individual cell is tracked under the action of the optics. Under the effect of self-adaption and levitation, a single cell can maintain a stable spatial position. Scale bar: 50 μm.

#### Single live-cell imaging tracking

Single-cell imaging is used to study cell heterogeneity in cancer line cases. However, direct monitoring of many individual initial cells with morphological change over time has so far not been technically feasible[90], [91]. Single cell tracking in 3D space is possible and combined with subsequent biochemical analyses of individually tracked cells, keeping their identity traceable with the CELLviewer system. Single-cell 2D tracking can more easily integrate subsequent biochemical analyses and act as surrogate measurements for the 3D situation[74]. On this basis, the 3D rotation of single cells was successfully achieved experimentally in figure Fig. 5.

### Generation of regular 3D spheroids for CELLviewer analysis

#### Materials

- *MCF-7* cell line (ATCC^®^ HTB-22TM)
- DMEM with 1 g/L glucose, sodium pyruvate and L-glutamine (Corning^®^ Life Sciences)
- FBS 10% (GibcoTM, Life Technologies, Thermo Fisher Scientific)
- L-glutamine 2 mM (Sigma-Aldrich, Merck)
- Penicillin-Streptomycin solution (Sigma-Aldrich, Merck)
- Dulbecco’s Phosphate Buffered Saline with MgCl2 and CaCl2 (Sigma-Aldrich, Merck)
- Percoll^®^ solution, pH 8.5-9.5 (25°C), cell culture tested (Sigma-Aldrich, Merck)
- Sphericalplate 5D^®^ 24-well cell culture plate (Kugelmeiers AG)
- MitoGreen (PromoKine, PromoCell)
- CELLviewer imaging system and disposable cartridge
- CELLviewer 50 mL DOCK
- ImageJ software (US National Institutes of Health)

#### Methods

*MCF-7* cells were grown at 37 °C and 5% CO2 in DMEM medium, supplemented with 2 mM L-glutamine, 10% FBS, 100 units/mL penicillin and 100 mg/mL streptomycin. Under a laminar flow sterile hood, the Sphericalplate 5D^®^ plate is rinsed first with 1 mL of sterile PBS followed by 1 mL of complete medium. Every well is preloaded with 0,5 mL of complete medium, then 0,5 mL of cell suspension at a concentration of 3×10^5^ cells/ mL is pipetted in every well for a total volume of 1 mL per well. The plate is then incubated at 37 °C and 5% CO2 for 24 hours to promote spontaneous cell aggregation and uniform-sized 3D spheroids formation. The 3D Spheroids are withdrawn from 4 wells and trasferred in a 50 mL centrifuge tube. Sample is centrifuged for 5 minutes at 800 rpm and the surnatant is discarded by gently aspirating. The pellet is resuspended in DMEM medium supplemented with Percoll^®^, to improve sample stable focusing during long-term imaging in CELLviewer cartridge. Basicly, 3000 spheroids are resuspended in 20 mL of Percoll^®^ supplemented DMEM medium to achieve a working concentration of 150 spheroids/mL. Consequently, MitoGreen 400nm solution is added to the culture medium and the sample is then pipetted inside a 50 mL Falcon tube closed with a CELLviewer 50 mL DOCK. MitoGreen probe uptake is thereby quantitatively analyzed within CELLviewer over the experiment lifetime. CELLviewer digitally isolates a single 3D spheroid and focuses the sample with a fluidic feedback mechanism. CELLviewer software, CELLcontrol, manages the imaging setup to automatically acquire sample images in Bright-field channel and GFP channel at 0,5 fps with 20X magnification. ImageJ software is used for image analysis using Measure function to calculate 3D spheroids diameter. All image acquisitions from FITC channel are stacked together and the same square shaped ROI (including spheroid borders) is applied to all the images. Adjust Brightness/Contrast function is used to homogeneously remove fluorescence background to all the stack images. Max Grey Value function is used to quantitatively assess fluorescence signal increase over the experiment lifetime.

#### Result

1,5×10^5^ *MCF-7* cells are seeded in 1 mL of complete medium for every well to let them aggregate. Homogenous-sized spheroid population shows mean diameter of 73 ± 4 μm at 24 hours of incubation and mean diameter of 73 ± 5 μm after 48 hours (Fig. 6).

**Fig. 6.**
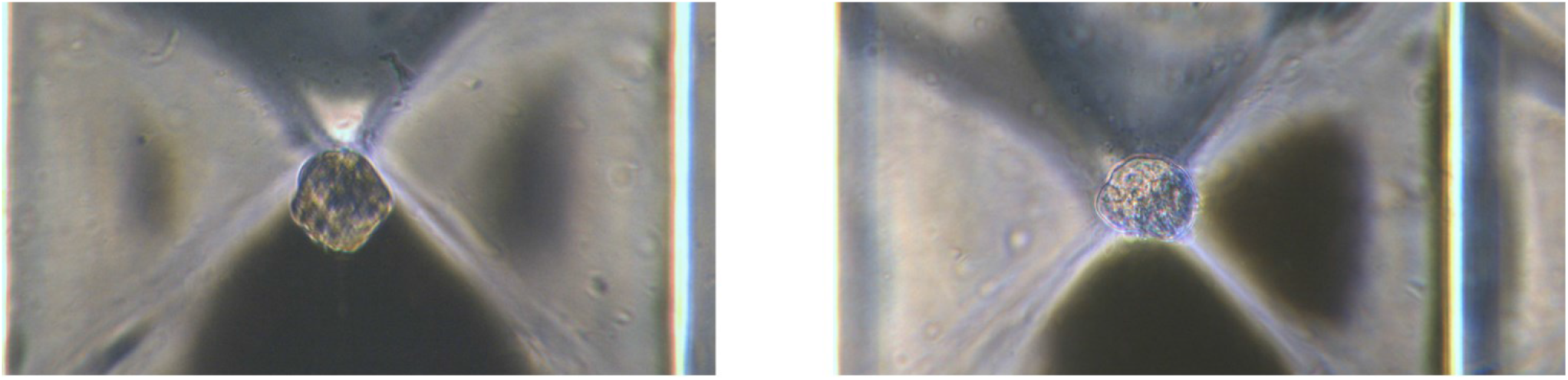
Bright-field analysis of *MCF-7* spheroids grown in Sphericalplate 5D^®^. Left: Bright-field acquisition at 24 hours. Right: Bright-field acquisition at 48 hours. Image acquisition with Leica Microsystems inverted Epi-fluorescence microscope DMLB Fluo MS15062. Scale bar: 50 μm.

Spheroids diameter remains constant within 24 and 48 hours. Single 3D spheroids, composed of different cell lines (depending on customers’ requirements) are digitally isolated in CELLviewer cartridge and a fluidic feedback mechanism focuses the sample in the analysis chamber for time-lapse, long-term culture and time-lapse live imaging. As shown in Figure 7 below, 100 μm diameter *MCF-7* spheroid is cultured for 4 hours in CELLviewer. A bright green fluorescent signal assesses spheroid viability, since MitoGreen conceptually overlaps mitochondrial regions of all viable cells composing a 3D spheroids. Fluorescence signals gradually increases throughout the experiment time course, reaching a maximum signal intensity within the first 1 hour and a half from the experiment start, due to MitoGreen probe progressive uptake by multicellular 3D spheroids.

**Fig. 7.**
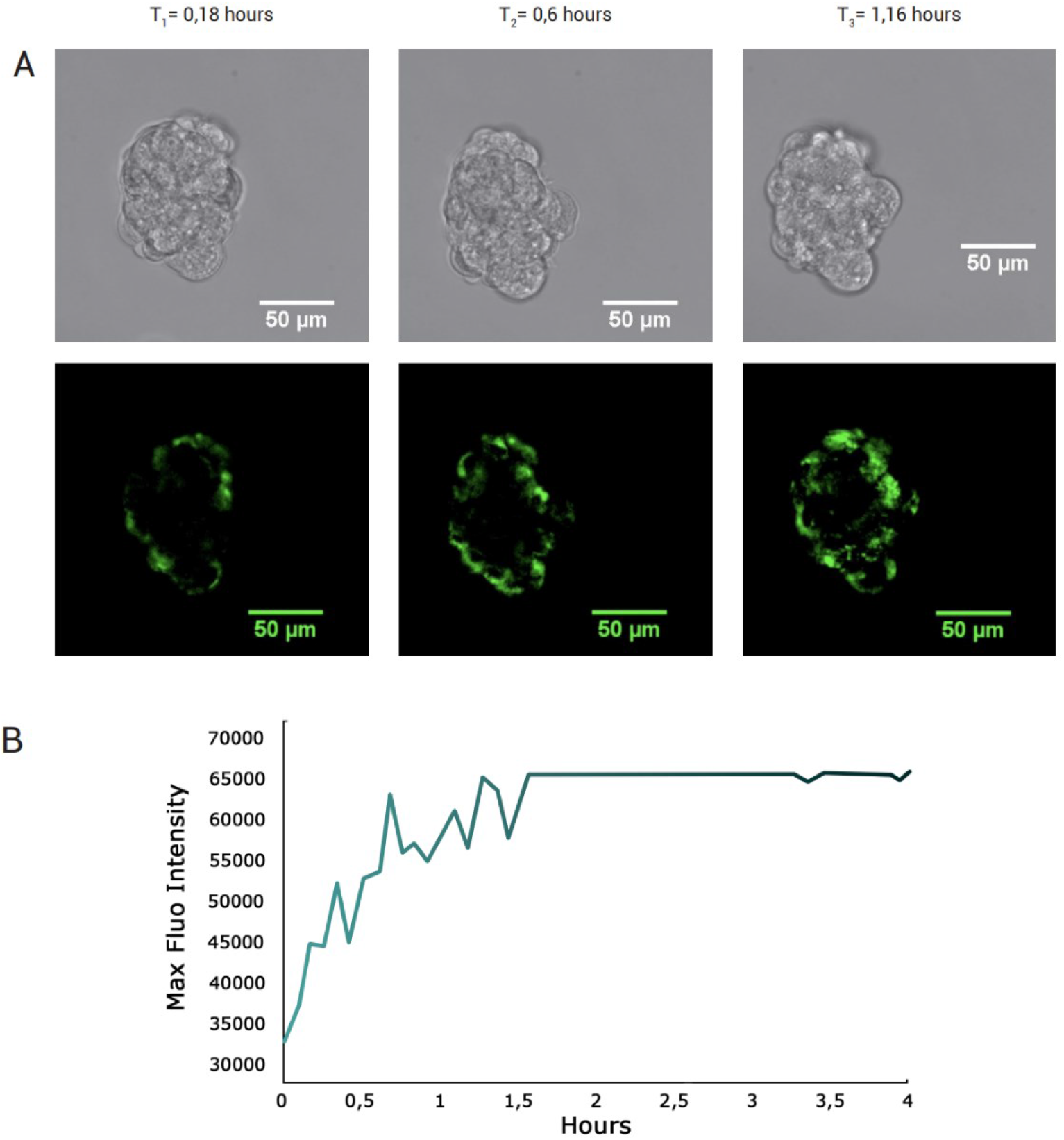
Analysis of MitoGreen labelled *MCF-7* spheroid. **A** CELLviewer acquisitions in Bright-field channel (top) and in GFP-channel (below) at 3 different timepoints. **B** Scatter Plot of MitoGreen Max Fluorescence Intensity (Grey Values) over the course of experiment. Scale bar: 50 μm.

## DISCUSSION

Compared to other devices in the literature[67], [72], [77], [84], [92] the CELLviewer is able in addition to obtaining detailed images of current cellular morphology with resolution and high-quality data; it is possible to carry out experiments in direct time in terms of: physiology, toxicology, and clinical pharmacology. The entire automated system allows full autonomy and protocol management thanks to the software making the operator free to conduct other work, thus increasing the productivity of his project. A microfluidic system has been developed and demonstrated that the 3D model can be spatially located the 3D model. The Epi-Fluorescence mitochondrial imaging was conducted to guide the configuration. Moreover, by utilizing time-lapse imaging of cells can be achieved, the evolution of cells and their 3D morphology can be acquired. In summary, the proposed microfluidic technology can serve as a new platform approach, which has the potential to advance studies at the cellular level. Given the basic importance of the research economy, the maximization of the times and ways of producing information is a close and obligatory landing. The preliminary experience conducted with CELLviewer indicates that this equipment responds to the need of individual operators as it consists of a synthesis of different integrated tools, which works both with manual and automated control. This kind of control system will facilitate the search dynamics even remotely, through software and on different devices, allowing the operator to simultaneously perform other activities that require his physical presence while ensuring maximum quality of the processes implemented. The live-imaging of 3D models in Microfluidics represents an optimal solution in observing the maintaining optimal health of cells while achieving the gap between the 2D model[93]. Recent advances in the design, prototyping, and production of microfluidic systems have enabled new ways of addressing disease study, with the advent of lab-on-a-chip technologies integrating different laboratory operations into individual microfluidic networks and advancing the understanding of cell-cell and cell-biomaterial interactions, with the engineering of organ-on-a-chip devices that mimic the response whole organs and systems using multichannel cell culture chips. These models are beginning to replace the most common cell culture systems, mainly Petri dishes, as the multichannel structure provides cells with a 2D or 3D-like environment to current *in vivo* configurations. Despite the progress of the last decade in the field of bioengineering, mainly concerning the success of these organ-on-chip systems as relevant research tools for the study of complex pathologies sustainably and economically, there is room for performance optimization. The development and use of devices capable of mimicking *in vivo* eukaryotic cells is a complex problem in itself, to understand the behavior and interactions with the extracellular environment capable of mimicking *in vivo* performance by advancing the research of the disease, constitutes a long-sought goal and an ever-present research challenge. Biomedical research on cellular systems of different complexities has evolved over the years towards 3D in which 2D has been deemed no longer enough.

The fields of biotechnological and biomedical research continually move towards a system that identifies as appropriate, a model capable of representing the *in vivo* counterpart, striving to validate new approaches with better results than the classic 2D monolayers in more physiological environments. Depending on the circumstances, there are different and often multiple reasons that differ between 2D and 3D where the cell cycle of a single cell alters or that the behavior of cell populations varies in space-time within clusters. The organization, composition, and several structures are among the best-understood signals that are integrated by the cell to regulate many key tasks including survival, differentiation, proliferation, migration, and polarization[94]. Inevitably, the study of biology and cell function *in vitro* requires stripping the cells of this native cell-cell and multi-cell interactions and their introduction into an environment of suspension and specific adhesion to the culture system. A better understanding of 3D model will help further optimize *in vitro* systems for the study of cell and tissue functions, as well as better translation to new therapeutic approaches.

## CONCLUSION

The main motivation for conducting experiments on 3D models also using the co-culture approach is to better mimic the target *in vivo*, study the evolution of the model and understand cell-cell interactions of any kind, both natural and engineered. These criteria are strongly influenced by the extracellular environment, susceptible to the configuration of the experimental design. Therefore, it is necessary to develop experimental systems capable of replicating the environment in which the 3D model will ultimately be used.

Staining of mitochondria with fluorescent dyes, antibodies or fluorescent molecules can greatly facilitate studies of their function and distribution and the viability of cells in healthy and diseased individuals. The preliminary experience conducted with CELLviewer indicates that this equipment responds to the needs of individual operators as it consists of a synthesis of different integrated tools, which works both with manual and automated control. A microfluidic system has been developed and demonstrated that the 3D model can locate the 3D model spatially, it’s possible to carry out experiments in direct time in terms of physiology, toxicology and clinical pharmacology.

The entire automated system allows full autonomy and protocol management thanks to the software making the operator free to conduct other work, thus increasing the productivity of his project. In summary, the proposed microfluidic technology can serve as a new platform approach, which has the potential to advance studies at the cellular level[95]. Fluorescence live-cell imaging is an indispensable tool for studying the structure and dynamics of cells and their internal/external building blocks[96].

Given the first considerations, the CellDynamics company is increasing the operating range of the CELLviewer machine, taking the cultivation of an even more complex 3D sample such as the zygote as a future approach. However, the main limitation of this machine turns out to be coupled with the complexity of that particular model: the speed of development and the maturation over time make it particularly difficult and unstable to manage with the system. These aspects need further investigation, but the gauntlet has been accepted.

## ACKNOWLEDGEMENTS

This article and the research behind it would not have been possible without the support of my supervisor Daniele Vigo. His need to train highly competent people, with the ability to grow independently, has been a source of inspiration since we had the first discussion outside. from class. His subject matter in the Veterinary Biotechnology course has opened my eyes to a world so vast and rich in knowledge, fascinating and engaging. For the examination of him I began to do research and have never stopped since. The professor has always been a support within the university and he immediately became interested in me like no other university professor. He has always encouraged me to hurry up to get a degree and work together, unfortunately I failed in this and I’m sorry to have disappointed him. Until the end, at least until now he has always been lenient. Not only by sharing his future projects but also his complete willingness to help me in the next steps.

I thank Simone Pasqua and the members of CellDynamics who have been able to enhance their projects and create a truly incredible company, making possible what I have described in the previous chapters. Not only has this experience with them increased my vision but it has left me with a void that I hope to fill with what awaits me.

I also thank Gabriele Brecchia for his unfailing patience and for his kindness to grant a place in his office to a shaven person as I am. I am grateful to him for taking the time to listen to my fears and for introducing me to the world of European Congresses, which I would never have been able to access without him. Finally, I thank you for having endured the stench of burning when I welded the various components of the bioreactor.

Finally, I thank Giulio Curone for his wisdom shared with me, little pearls that have not been lost. A person who has long vision, it is no coincidence that you are Professor Vigo’s favorite. I believe that without you it would not be the same. Like all the other people mentioned, I admire you for your commitment and wish you the best.

Finally, I thank my parents, I have no words for what I would like to write. Facts matter more than words and I want it to be so. Without them I would not have achieved anything, and their sacrifices will not be in vain, in the hope of being able to fulfill myself in life.

A hug to everyone, thank you for your presence on this path.

